# PhISCS-BnB: A Fast Branch and Bound Algorithm for the Perfect Tumor Phylogeny Reconstruction Problem

**DOI:** 10.1101/2020.02.06.938043

**Authors:** Erfan Sadeqi Azer, Farid Rashidi Mehrabadi, Xuan Cindy Li, Salem Malikić, Alejandro A. Schäffer, E. Michael Gertz, Chi-Ping Day, Eva Pérez-Guijarro, Kerrie Marie, Maxwell P. Lee, Glenn Merlino, Funda Ergun, S. Cenk Sahinalp

**Affiliations:** Department of Computer Science, Indiana University, Bloomington, IN 47408, USA; Program in Computational Biology, Bioinformatics, and Genomics, University of Maryland, College Park, MD 20742, USA; Cancer Data Science Laboratory, Center for Cancer Research, National Cancer Institute, National Institutes of Health, Bethesda, MD 20892, USA; Laboratory of Cancer Biology and Genetics, Center for Cancer Research, National Cancer Institute, National Institutes of Health, Bethesda, MD 20892, USA

**Author notes:** Joint first authors (Erfan Sadeqi Azer, Farid Rashidi Mehrabadi).

## Abstract

**Motivation:** Recent advances in single cell sequencing (SCS) offer an unprecedented insight into tumor emergence and evolution. Principled approaches to tumor phylogeny reconstruction via SCS data are typically based on general computational methods for solving an integer linear program (ILP), or a constraint satisfaction program (CSP), which, although guaranteeing convergence to the most likely solution, are very slow. Others based on Monte Carlo Markov Chain (MCMC) or alternative heuristics not only offer no such guarantee, but also are not faster in practice. As a result, novel methods that can scale up to handle the size and noise characteristics of emerging SCS data are highly desirable to fully utilize this technology.

**Results:** We introduce PhISCS-BnB, a Branch and Bound algorithm to compute the most likely perfect phylogeny (PP) on an input genotype matrix extracted from a SCS data set. PhISCS-BnB not only offers an optimality guarantee, but is also 10 to 100 times faster than the best available methods on simulated tumor SCS data. We also applied PhISCS-BnB on a large melanoma data set derived from the sub-lineages of a cell line involving 24 clones with 3574 mutations, which returned the optimal tumor phylogeny in less than 2 hours. The resulting phylogeny also agrees with bulk exome sequencing data obtained from *in vivo* tumors growing out from the same cell line.

**Availability:** https://github.com/algo-cancer/PhISCS-BnB

## 1 Introduction

Cancer is a highly dynamic, evolutionary disease. Constantly shaped by mutation and selection, cancer progression often results in the emergence of distinct tumor cell populations with varying sets of somatic mutations, commonly known as (sub)clones. The diverse pool of subclones may harbor treatment-resistant mutations. When favorably selected for in the tumor environment by treatment exposure, treatment-resistant subclones may gain dominance over others and eventually contribute to treatment failure (Alizadeh et al., 2015). The challenges in developing effective cancer therapies under the heterogeneous tumor landscape thus motivate the following question: can we reconstruct the tumor phylogeny and unravel spatial and temporal intra-tumor heterogeneity (ITH) to enlighten cancer treatment strategies?

In recent years, several computational tools for analyzing intra-tumor heterogeneity and evolution from bulk sequencing data of tumor samples have been developed (Strino et al., 2013; Jiao et al., 2014; Hajirasouliha et al., 2014; Deshwar et al., 2015; Popic et al., 2015; El-Kebir et al., 2015; Malikic et al., 2015; Marass et al., 2016; El-Kebir et al., 2016; Donmez et al., 2017; Satas and Raphael, 2017). However, bulk sequencing data provides only an aggregate signal over large number of cells and, due to its limited resolution, all of these methods have several limitations in unambiguously inferring trees of tumor evolution. Most notably, they typically rely on clustering of mutations of similar cellular prevalence. Consequently, if two sets of mutations evolving on different branches of phylogenetic tree have similar cellular prevalence values, they get clustered together. Furthermore, even in cases where the cellular prevalence values of clusters are different, methods based on bulk sequencing data are frequently unable to distinguish between multiple trees that describe the observed data equally well (Malikic et al., 2019a; Kuipers et al., 2017).

The rise of single-cell sequencing (SCS) has enabled exploration of ITH at a higher, cellular resolution. Unfortunately even SCS can not trivially provide a comprehensive understanding of ITH. Among the lingering caveats with SCS, the most prominent is the prevailing presence of sequencing noise (Zafar et al., 2018). We are particularly interested in three types of noise in SCS datasets. (1) False positive mutation calls, potentially from sources like read errors, (2) false negative mutation calls, potentially from sources like variance in sequence coverage or allele dropout, (3) missing values for mutations from sites affected by DNA amplification failure.^1^ The multi-faceted and high levels of sequencing noise have prompted the development of novel computational approaches that need to infer a tumor evolutionary model while compensating for all three sources of noise.

The first principled approaches for studying ITH by the use of SCS data were all based on probabilistic formulations that aim to infer the *most-likely perfect-phylogeny* (PP) of a tumor. SCITE (Jahn et al., 2016), OncoNEM (Ross and Markowetz, 2016) and SiFit (Zafar et al., 2017) are among these methods that primarily aim to build a PP (i.e. an evolutionary tree where no mutation can appear more than once and is never lost).^2^ Following up on this, SPhyR (El-Kebir, 2018) formulates the tumor phylogeny reconstruction problem as an integer linear program (ILP) under the constraints imposed by the *k*-Dollo parsimony model - where a gained mutation can only be lost *k* times. SCIΦ (Singer et al., 2018). SPhyR simultaneously performs mutation calling and the tumor phylogeny inference taking read counts data as the input, rather than the more commonly used genotype matrix with inferred mutations, represented by columns, in distinct cells, represented in by rows.

As datasets with matching SCS and bulk sequencing data become publicly available, methods to infer tumor phylogeny through joint use of these two data types are becoming available. B-SCITE (Malikic et al., 2019a) for example combines CITUP, which is designed for bulk sequencing data, with SCITE through an MCMC strategy. More recently PhISCS (Malikic et al., 2019b), offers the option of formulating integrative reconstruction of the most likely tumor phylogeny either as an ILP or as a CSP (boolean constraint satisfaction program), while allowing for a fixed number of PP violating mutations. The CSP version of PhISCS, when employing state of the art CSP (more specifically weighted max-SAT) solvers such as RC2 (Ignatiev et al., 2019) and Open-WBO (Martins et al., 2014) turn out to be the fastest among all available techniques even when only SCS data is available. Nevertheless, none of the available techniques can scale up to handle emerging data sets that involve thousands of cells (Laks et al., 2019); even moderate size SCS data involving a few hundred mutations and cells turn out to be problematic especially with the current “standard” false negative rate of 15 − 20%. Other recent techniques such as scVILP (Edrisi et al., 2019) and SiCloneFit (Zafar et al., 2019) focus on adding new features to (or relax constraints for) the tumor phylogeny reconstruction problem and are (typically) not faster.^3^ Finally, even though new sequencing techniques such as single “clone” sequencing (SClS, i.e. bulk sequencing of homogeneous cell populations, each derived from a single cell) offer much lower false negative rates, the scale of the data they produce - involving thousands of mutations, require much faster solutions to the tumor phylogeny reconstruction problem.

In this paper, we present a Branch and Bound (BnB) algorithm and its implementation (called PhISCS-BnB, <u>Ph</u>ylogeny Inference using <u>S</u>ingle <u>C</u>ell <u>S</u>equencing via <u>B</u>ranch a<u>n</u>d <u>B</u>ound) to optimally reconstruct a tumor phylogeny very efficiently. Generally speaking, our BnB approach clusters entries of the input genotype matrix and processes them together, enabling faster execution. (See Section 3.1 and lemma 3.1). We introduce a number of *bounding* algorithms, some faster but offering limited pruning and others slower but with better pruning efficiency. Among them, a novel bounding algorithm with a 2-SAT formulation is a key technical contribution of our paper: through its use, PhISCS-BnB improves the running time of the fastest available methods for tumor phylogeny reconstruction by a factor of up to 100. (See Section 4).

## 2 Perfect Phylogeny Reconstruction Problem

Given a binary genotype matrix *I*, we would like to reconstruct the most likely phylogeny by discovering how to flip the smallest number of entries of *I* so it can provide a Perfect Phylogeny.

### Preliminaries

Our input is a binary (genotype) matrix *I* ∈ {0, 1}^*n,m*^. The *n* rows represent genotypes of single cells observed in a single-cell sequencing experiment and the *m* columns represent a set of considered mutations. *I*(*i, j*) = 1 indicates that mutation *j* is present in cell *i*; *I*(*i, j*) = 0 indicates that it is not.

The *three-gametes rule* stipulates that a binary matrix *X* ∈ {0, 1}^*n,m*^ should *n*ot have three rows and two columns (in any order) with the corresponding six entries displaying the configuration (1, 0), (0, 1) and (1, 1). If the forbidden configuration is present, we say that there is a *violation*, referenced by the three rows and the two columns containing it. It was shown in (Gusfield, 1991) that satisfaction of the three-gametes rule by *I* is necessary and sufficient for the existence of a Perfect Phylogeny (PP) corresponding to *I*.

Given input matrix *I*, we call a binary matrix *X* a *descendant* of *I* if all entries of *X* are identical to those of *I* except some that have been *flipped* from 0 to 1. For a matrix *X, F*_0→1_(*I, X*) is defined as the number of entries that are 0 in *I* and 1 in *X*. We sometimes refer to this value as the number of flips to get to *X* from *I*.

### Our Problem

Given a genotype matrix *I*, we would like to obtain a minimum-cardinality set of *bit flips* (from 0 to 1) ^4^ that removes all three-gametes rule violations in *I* and thus transforms *I* into a matrix *Y* that provides a PP.

## 3 Branch and Bound Method

In order to discover the smallest number of 0 to 1 flips that will remove all violations in input matrix *I*, we use a branch and bound (BnB) technique. In what follows, we give an overview of the building blocks of our branch and bound approach. Then we put all of them together in Algorithm PhISCS-BnB.

Our Branch and Bound algorithm forms a search tree where each node contains a matrix, with input matrix *I* at the root – for simplicity we might refer to a node with its label as well as its matrix. In this tree, a matrix *X* at node *v* is a *descendant* (as described in the preliminaries) of the matrix *Y* at *v*’s parent node; all matrices in the tree are thus descendants of *I*. The tree terminates in leaf nodes that are PP; non-PP nodes will have two child nodes as the tree grows unless they have been pruned due to detected nonoptimality.

When a node *v* with matrix *X* is formed, *v* is assigned a *priority score* equal to the number of bit flips needed to get from *I* to *X* plus a lower bound on the number of flips necessary to remove all the violations in *X*. All nodes are kept in a priority queue and are explored in ascending order of their priority scores, unless they have been removed from consideration (pruned) by the bounding mechanism. When the whole tree has been explored or pruned, one of the PP nodes with the smallest number of flips away from *I* yields the answer.

For matrix *X*, we let 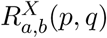 denote the set of rows with *a* in column *p* and *b* in column *q*, i.e., 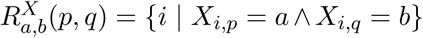. We drop the superscript, when the matrix *X* is clear from the context, and only write *R*_*a,b*_(*p, q*).

### AAlgorithm 1 PhISCS-BnB

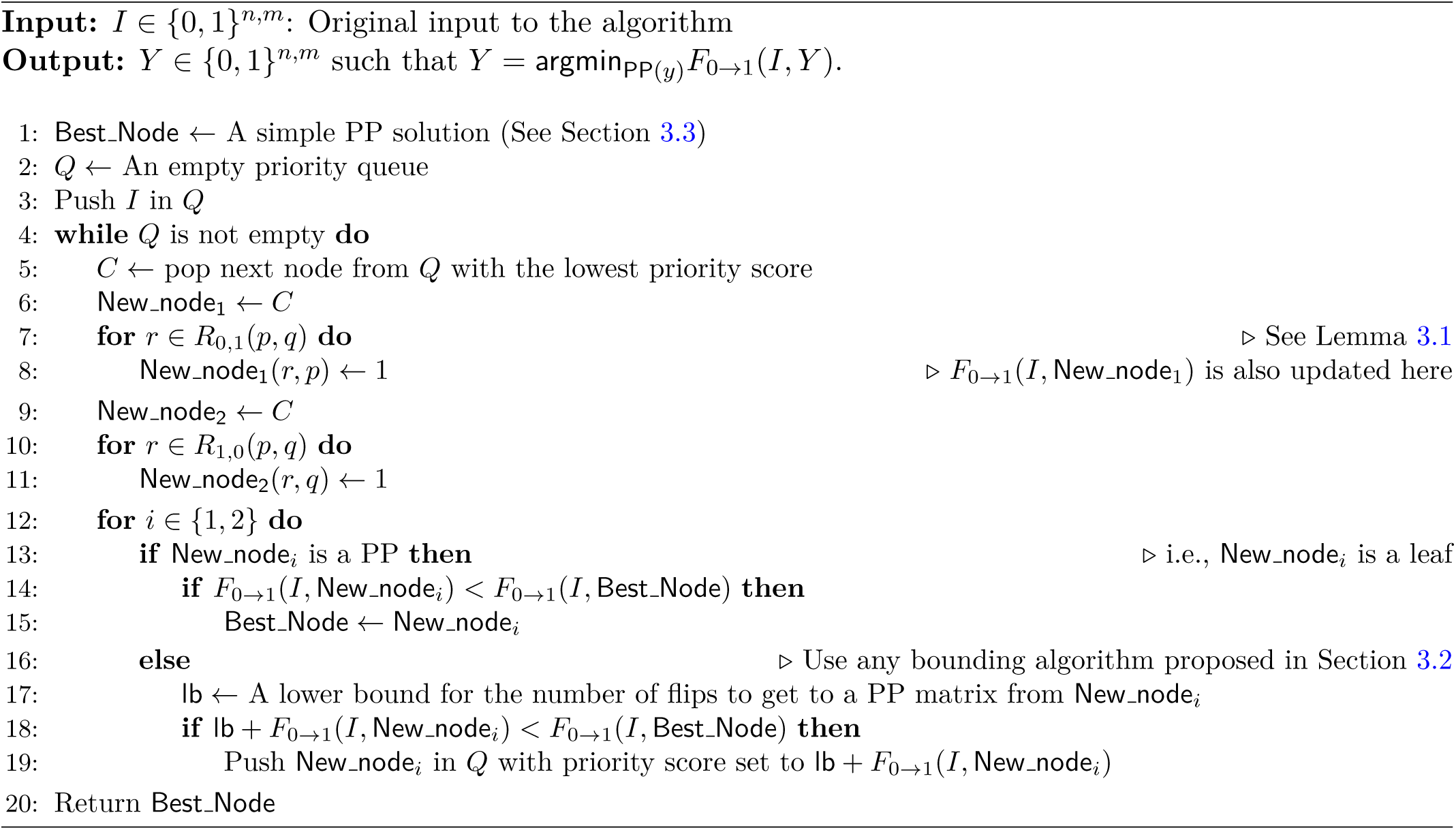

### 3.1 Branching

Let *X* be the matrix at the node being explored. If *X* has no violation, it is considered a leaf. Otherwise, let (*p, q*) be the pair of columns for one particular violation that was found,^5^ i.e., |*R*_*a,b*_(*p, q*)| > 0 for all (*a, b*) ∈ {(0, 1), (1, 0), (1, 1)}. We have two options for fixing the violation. As a violation involving columns *p, q* contains both a (1, 0) and a (0, 1) in different rows, we have the option of converting either one to a (1, 1) to remove the violation. To reflect this, we construct two child nodes from the current node, one for each option. As an added optimization, once we decide to fix a (0, 1) (resp. (1, 0)) on columns *p, q*, we fix all (0, 1) (resp. (1, 0)) on these two columns, by changing them to (1, 1). In particular, in the left child, all entries whose row is in *R*_0,1_(*p, q*) and whose column is *p* are flipped from 0 to 1. Similarly, in the right child entries whose row is in *R*_1,0_(*p, q*) and whose column is *q* are flipped.

In some cases, the above branching rule, which can flip multiple 0s at a time, shrinks the height of the search tree compared to the algorithm in (Chen et al., 2006; Cai, 1996), which flips a single 0 in a child node. The following lemma formally expresses why we flip several entries in a column together at Lines 8 and 11 of the pseudocode: if a (0, 1) in a violation involving columns *p, q* of matrix *X* is a (1, 1) in a PP descendant *X*′ of *X*, all other (0, 1) on (*p, q*) are (1, 1) as well. An analogous statement holds for (1, 0).

#### Lemma 3.1.

*For any X* ∈ {0, 1}^*n,m*^ *with a violation involving columns p, q, let X*′ *be any PP descendant of X. Then, at least one of the following hold:*

- ∀*r*_1_ ∈ *R*_0,1_(*p, q*), *X*′(*r*_1_, *p*) = 1, *or*
- ∀*r*_2_ ∈ *R*_1,0_(*p, q*), *X*′(*r*_2_, *q*) = 1.

*Proof.* Assume that the lemma is false; i.e., there is an *X*′ such that ∃*r*_1_ ∈ *R*_0,1_(*p, q*), *X*′(*r*_1_, *p*) = 0 ∧ ∃*r*_2_ ∈ *R*_1,0_(*p, q*), *X*′(*r*_2_, *q*) = 0.

Since (*p, q*) corresponds to a violation and all the 1 entries in *X* have to remain 1 in *X*′, there should be a row *r*_3_ such that *X*′(*r*_3_, *p*) = *X*′(*r*_3_, *q*) = 1. This implies that the pair of columns (*p, q*) and the triplet of rows (*r*_1_, *r*_2_, *r*_3_) corresponds to a violation. This contradicts the assumption that *X*′ is PP. □

### 3.2 Bounding Mechanisms

A *bounding* algorithm is a method that computes a *l*ower bound for the number of flips needed to transform matrix *X* at a node *v* to a PP matrix. It thus helps the branch and bound algorithm to prune the nodes that are provably worse than the currently maintained best node, i.e., the variable Best Node in Algorithm PhISCS-BnB.

We reuse the calculated lower bound as the estimate of how many flips a matrix *X* will require to transform into a PP matrix, and then add the number of flips needed to transfer *I* to *X*, in order to set the priority score of the node containing *X*. Recall that all the introduced nodes are pushed to a priority queue, and in each iteration the node with the lowest priority score is chosen to be explored. So the node *X* we pick will represent the lowest number of total flips from *I* to a PP node going through *X*.

One observation that leads to a lower bound is as follows. Consider a pair of columns that have at least one row with pattern (1, 1), then the number of flips, involving columns *p* and *q*, is at least min(|*R*_0,1_(*p, q*)|, |*R*_1,0_(*p, q*)|). Therefore, for an arbitrary partitioning of the set of columns to pairs, we can aggregate these bounds to achieve a lower bound for the whole matrix.

In the above, one would expect the choice of the partition to have an impact on the quality of the lower bound estimate. To explore this, in what follows, we present three bounding algorithms that progressively add more sophistication to the above idea. These three bounding algorithms offer a tradeoff between the per-node running time and the accuracy of the bound. For some inputs the fast (and possibly not-so accurate) bounding results in a faster execution, but for other inputs a different tradeoff is better. It is worth mentioning that for biologically plausible inputs, our experiments show that the higher accuracy of bounding is much more important to total time than the per-node running time. We present first two bounding algorithms in Section 3.2.1. The third and the most sophisticated one is presented in Section 3.2.2 and is used in our experiments to compare with previous tools in the literature.

#### 3.2.1 Random Partition vs Maximum Weighted Matching

As our first bounding method, we partition the columns of the matrix into pairs uniformly at random. The technique is simple, but more sophisticated techniques might give tighter bounds.

As our second method, we describe a method based on Maximum Weighted Matching (MWM). Construct a weighted undirected graph *G* = (*V, E, w*) where the vertices are the columns of *I*, each column representing a mutation: *V* = {*c*_1_, … *c*_*m*_} (*c*_*i*_ corresponds to *i*^*th*^ mutation) and the edges are column pairs that display a (1, 1): *E* = {{*c*_*i*_, *c*_*j*_} | |*R*_1,1_(*c*_*i*_, *c*_*j*_)| > 0. In the following we calculate a weight corresponding to an edge *e* = {*c*_*i*_, *c*_*j*_} ∈ *E*, to be a lower bound on the number of entries in columns *c*_*i*_ and *c*_*j*_ that have to be flipped to make *I* a PP matrix. Formally, for each edge *e* = {*c*_*i*_, *c*_*j*_} ∈ *E, w*(*e*) = min(|*R*_0,1_(*c*_*i*_, *c*_*j*_)|, |*R*_1,0_(*c*_*i*_, *c*_*j*_)|). The process of constructing *G* takes Θ(*nm*^2^) time. In this graph theoretic formulation, each partitioning corresponds to a matching *G*. Thus, we take advantage of the algorithm described in (Galil, 1986) to find maximum weighted matching with *O*(*m*^3^) running time.

In both bounding algorithms, one can maintain the bounds dynamically by processing only small changes from one node to another node near it, (in some cases, to its sibling). The details of such dynamic maintenance are out of the scope of this work.

#### 3.2.2 2-SAT

For this approach we present a novel constraint satisfaction formulation that describes a set, containing but not necessarily equal to, all the valid flips-set corresponding to *I*. Let *z*_*i,j*_ denote a binary variable corresponding to the (*i, j*)^*th*^ entry. We define variables for only zero entries. Consider a pair of columns (*p, q*) with |*R*_1,1_(*p, q*)| > 0. For any row *r*_1_ ∈ *R*_0,1_(*p, q*) and any row *r*_2_ ∈ *R*_1,0_(*p, q*) add 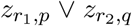. The intuition is that for any violation sextuplet, either one of zeros should be flipped. Satisfying these constraints is necessary to achieve a PP matrix, but not sufficient.

Let MWS (short for Minimum Weighted SAT) denote an arbitrary off-the-shelf tool that, given a satisfiable Boolean formula, outputs a satisfying assignment with the minimum number of variables assigned to true. Then the number of variables with true value in an optimal assignment, satisfying all these constraints, is a lower bound for the optimal number of flips resulting in a PP matrix. Formally, the lower bound is equal to

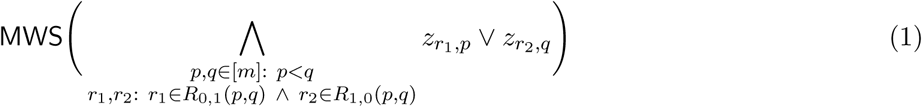

After achieving a minimum weight satisfying assignment, we flip those zero entries that correspond to *z* variables with value 1.

##### Compact Formulation

The formulation in Equation 1 can be expressed in fewer constraints by introducing a new set of variables and following the case distinction in Lemma 3.1: for each pair of columns *p, q* define a corresponding binary variable *B*_*p,q*_. The weight of this new set of variables is set to zero in MWS formulation. If this variable is set to zero (by a minimum weighted SAT routine) then all variables 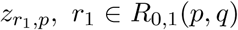, take value 1. Similarly, if the variable is set to one, then all 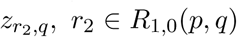 take value 1. Formally, the formulation changes to MWS(*H*_1_ ∧ *H*_2_), where,

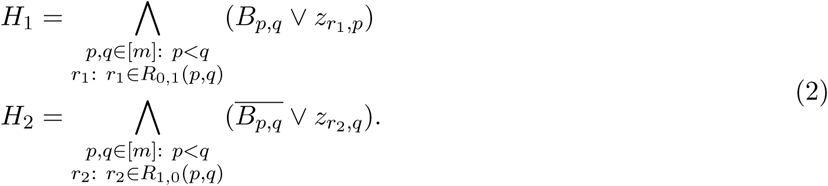

The number of constraints corresponding to the column pair (*p, q*) will decrease from |*R*_0,1_(*p, q*) | · |*R*_1,0_(*p, q*)| in Equation 1 to |*R*_0,1_(*p, q*)| **+** |*R*_1,0_(*p, q*)| in Equation 2. There are two advantages to the compact formulation: (I) the time spent on forming the set of constraints is shorter, (II) for some set of inputs, heuristic sat-solvers run more efficiently on the formulation given in Equation 2 than on Equation 1, even though they are logically equivalent. In each experiment, we use only one of these formulations and it is specified in the corresponding description of the experiment.

##### Extra constraints

As another version of our lower bound, we add a set of new constraints. These constraints improve the lower bound to be a closer estimate of the optimal number of flips for some inputs. The tighter bound helps the branch and bound framework explore fewer nodes, even though the time to compute the bound per node increases. This advantage comes with a dip in the running time of the bounding calculation within each node.

The idea for the new set of constraints is to preclude some solutions that satisfy all constraints in Equation 1 but still do not remove all violations. In particular, when, for a specific pair of columns (*p, q*), *R*_1,1_(*p, q*) is empty, there is no constraint involving pair (*p, q*) in Equation 1. As an example, assume Columns 1, 2 contain both (1, 0) and (0, 1) rows, but no (1, 1). Columns 2, 3 contain a violation, which MWS removes by flipping 0s in column 2. This might create a (1, 1) in Columns 1, 2, and create a violation that was not there in the beginning. Since, Equation 1 does not contain any constraints for Columns 1, 2, this new violation is not removed.

In order to avoid such an outcome, we add new constraints to our formulation. Now, if some flip introduces a new violation, the extra constraints will enforce at least one additional flip to remove the newly created violation(s). Formally, the proposed set of constraints to add to Equation 1 is *E*_1_ ∧ *E*_2_, where,

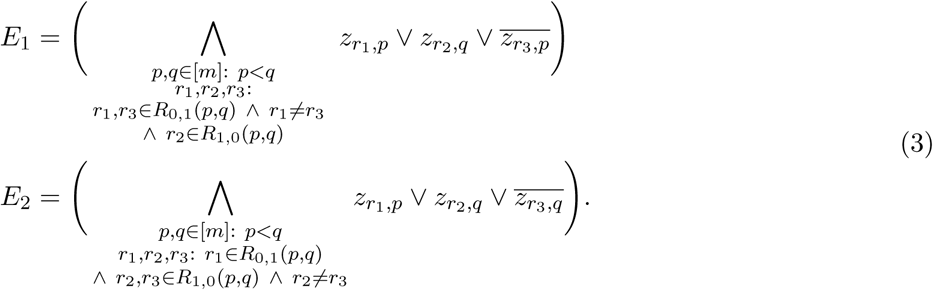

The number of constraints corresponding to columns (*p, q*) is |*R*_0,1_(*p, q*)| · |*R*_1,0_(*p, q*) | · (|*R*_0,1_(*p, q*)| **+** |*R*_1,0_(*p, q*)|). This is higher than the number of constraints in both Equations 1 and 2. However, for some matrices (e.g., the one processed in Section 4.1) the resulting tighter bound improves the running time of overall branch and bound algorithm tremendously.

### 3.3 Initial Solution

For the above bounding mechanism to start pruning, a feasible solution is required to initialize the variable Best Node at pseudocode Line 1. When using Random Partition or Maximum Weighted Matching as a bounding algorithm, find an initial value as follows. We first find a pair of columns corresponding to a violation and flip one of the zero entries involved in the violation. We repeat this until no violation is left. On the other hand, when 2-SAT bounding is used, we solve the corresponding formulation from Equations 1, 2, or 3. We then apply the chosen entries to flip and repeat this process until we obtain a PP matrix. In each iteration, at least one flip will be performed and there are finitely many zero entries to flip. Therefore, this process always terminates and results in a PP matrix.

### 3.4 Analysis

#### Correctness

In the search tree explored by the branch and bound algorithm, we are guaranteed to find the optimum path from *I* to a PP matrix. This is because throughout the execution (a lower bound on) the projected number of flips that a node needs to reach PP is compared against the currently best known way of reaching PP. If the node has no chance of beating the current best, it and all of its descendants are pruned. Consequently, PhISCS-BnB does not prune any nodes on the path to an optimum solution before reaching an optimum solution for the first time. Therefore, as long as our lower bounding techniques work correctly, our algorithm will discover an optimum PP.

In both the Random Partition and Maximum Weighted Matching techniques for obtaining a lower bound, we consider a partitioning of columns to pairs and calculate the minimum number of flips within each pair. Since the absence of violations within these pairs is necessary (but possibly not sufficient) for the removal of all violations, this estimate is a lower bound for the whole matrix.

In the 2-SAT approach, we form a set of constraints that must be satisfied in order to reach any PP descendant of a given node. That means the set of potential solutions obtained by satisfying all the constraints in the 2-SAT formulation is a superset of all PP matrices. That is why the optimum solution satisfying these conditions requires the same number of or fewer flips than any PP matrix.

#### Running Time

The worst-case running time of Algorithm PhISCS-BnB is 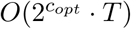 where *c*_*opt*_ is the minimum number of flips needed to turn *I* to a PP matrix, and *T* is the running time of the bounding algorithm. A naive bounding algorithm, as in (Chen et al., 2006; Cai, 1996), incurs a running time of *T* = *O*(*mn*). This is asymptotically the same as the running time for our random partition technique. Similarly, maximum weighted matching runs in time *T* = *O*(*nm*^2^ + *m*^3^): the first term is for forming the weighted graph, and the second term is for solving maximum weighted matching. Solving the 2-SAT formulation takes super-polynomial time in the worst case as the weighted 2-SAT problem is NP-hard. While one might expect infeasibly long running times due to the hardness of 2-SAT, our experiments generally seemed to avoid worst-case behavior as the running times stayed mostly reasonable even for large matrices.

## 4 Experimental Results

In this section we discuss our experimental results on real data and simulations.

### 4.1 Single Clone Sequencing Data from Mouse Melanoma Models

We first demonstrate the utility of PhISCS-BnB in studying evolutionary history on a large SClS data which contains a large number of mutations. The data set features the B2905 cell line and its sub-lineages derived from the “M4” mouse model of melanoma (Pérez-Guijarro et al., 2019). In brief, the Hgf-transgenic C57BL/6 pups received UV irradiation at postnatal day 3 (Zaidi et al., 2011). In 6-8 months, melanoma occurred in a fraction of UV-irradiated mice, and one of the harvested tumors was plated in culture to derive B2905 cell line (Pérez-Guijarro et al., 2019; Patel et al., 2017). To generate single-cell derived sub-lines, B2905 cells were harvested from *in vitro* culture, and subjected to fluorescence-assisted cell sorting (FACS) for isolating single cells in individual wells of 96-well plate. After expansion in culture, twenty four B2905 sub-lines derived from single cells were obtained. They are labeled C1 to C24.

After read alignment, calling and filtering the variants of all twenty four sub-lineages, 3574 distinct SNVs were detected in this dataset (details of these data analysis steps can be found in Supplementary File Section B). The clonal phylogeny depicted in Figure 1 was obtained by PhISCS-BnB in about 2 hours (we used the set of additional constraints mentioned in Section 3.2.2). The optimal solution obtained by PhISCS-BnB included 886 false negatives that were detected and corrected for establishing the mentioned phylogeny; this implies that the allele drop-out rate of the SClS protocol was around 1% as can be expected from bulk sequencing technology. For performance comparison purposes, we also ran the fastest available algorithmic tool (to the best of our knowledge), the CSP implementation of PhISCS (Malikic et al., 2019b) (namely PhISCS-B) on this data set, which required about 20 hours to obtain the same tree using the same computational platform (details of the computational platform can be found in Supplementary File Section C.2). On the other hand, the well known tool SCITE (Jahn et al., 2016), which is based on MCMC, could not report a result within approximately 24 hours of running.

**Figure 1:**
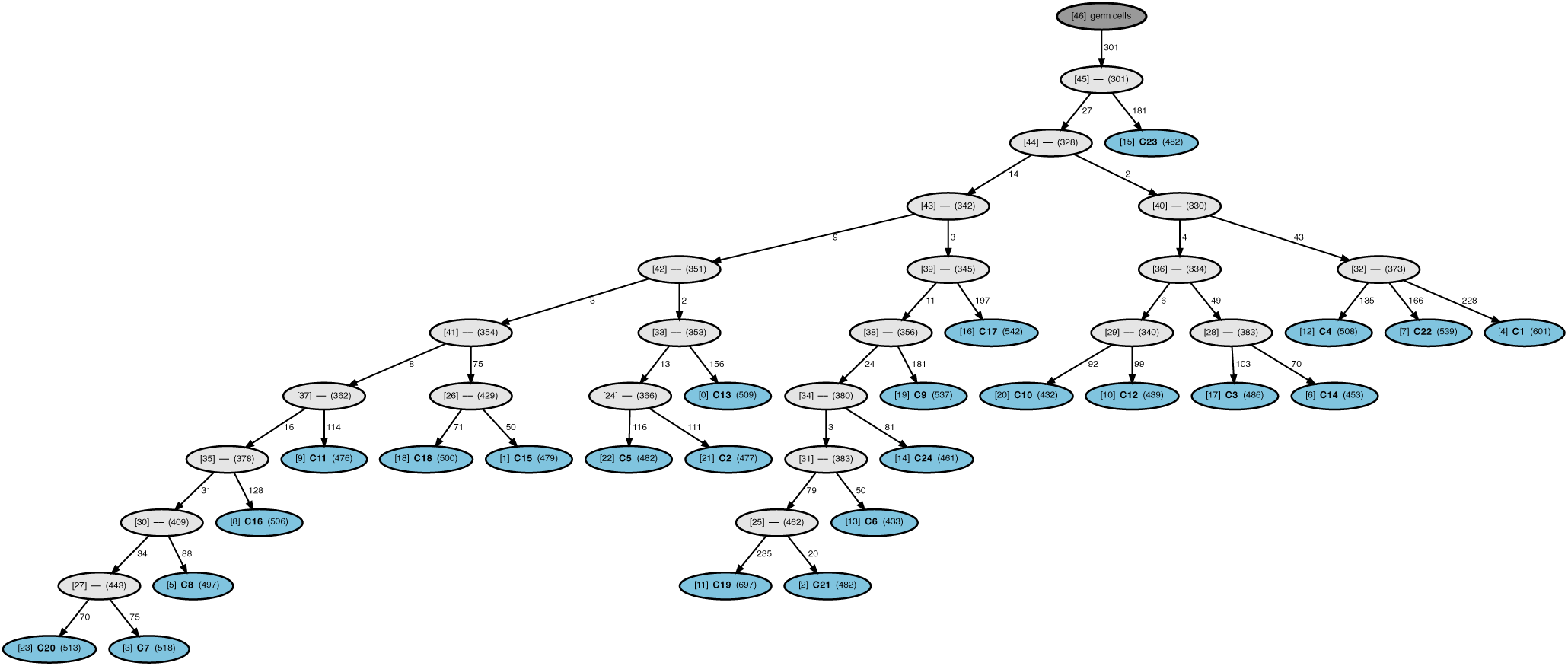
The clonal tree obtained by PhISCS-BnB from 24 clonal sub-lineages of B2905 cell line that are derived from the “M4” mouse model for melanoma. For each node, the number inside the brackets denotes its node id and the number inside the parentheses shows the total number of mutations occurring on the path from the germline (root) to the node (i.e., the total number of mutations harbored by the node). The edge labels represent the number of mutations occurring between a parent and its child node. The complete list of mutations occurring at each edge can be found at https://github.com/algo-cancer/PhISCS-BnB/blob/master/real/real.mutsAtEdges. The leaf nodes (colored blue) also include their sub-lineage labels.

We note that in an earlier study, the parental B2905 cell line was implanted to syngeneic mice to grow tumors. The mutations of these *in vivo* tumors were then identified via whole exome sequencing. Interestingly, only mutations associated with nodes 45, 44, 43, 42, and 41 in the Figure 1 were expanded *in vivo* when the parental line was implanted into syngeneic immunocompetent mice, suggesting that subclones associated with node 41 (C1, C14, C22, C16, C8, C7 and C20) survived better while others declined. Although the interpretation is limited by the small number (24) of subclones sampled from the parental cell line, it is consistent with the concept of *immunoediting* (Mittal et al., 2014) and implies the node-associated mutations may serve as markers to track dynamics and evolution of subclones in tumors. Moreover, the hierarchy of the mutations may help to delineate driver and passenger mutations (Schwartz and Schäffer, 2017).

### 4.2 Comparison of PhISCS-BnB against PhISCS-B and PhISCS-I on simulated data

Next, we compared the running time of PhISCS-BnB on simulated data against the fastest available algorithmic tool, PhISCS (Malikic et al., 2019b), which has two versions: PhISCS-B is based on CSP whereas PhISCS-I is based on ILP. Both were compared to our tool on simulated SCS data with 100 to 300 cells and 100 to 300 mutations, with false negative error rates ranging from 5% to 20%. In each case, 10 distinct trees of tumor evolution were simulated, each with 10 subclones. We allowed all three programs to run up to 8 hours on each simulated dataset (details of the computational platform we used can be found in Supplementary File Section C.1). Figure 2 clearly shows that PhISCS-BnB is faster than the best available alternative (i.e. PhISCS-B) by a factor of 10 to 100.^6^

**Figure 2:**
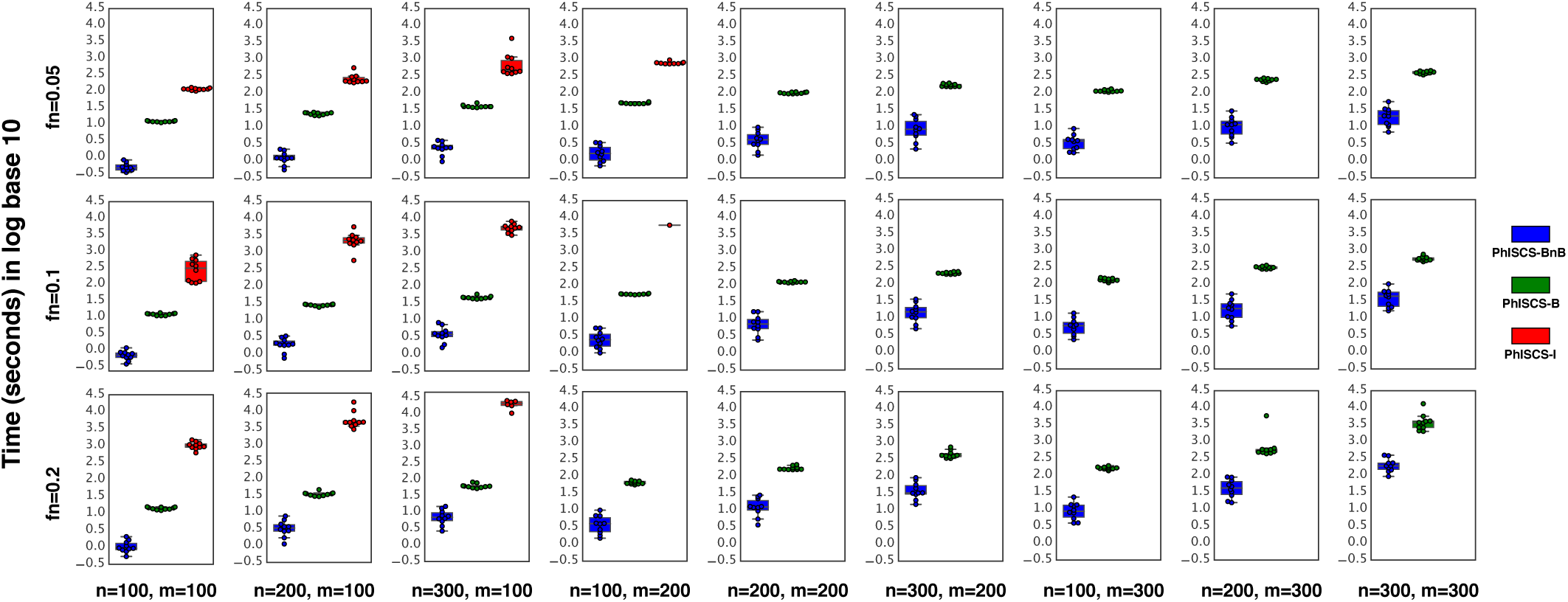
Comparison of PhISCS-BnB with PhISCS-B and PhISCS-I in terms of running time (seconds) in log base 10. In each case, 10 distinct trees of tumor evolution were generated, each with 10 subclones. A timelimit of 8 hours was used for running each tool (those cases that exceed the time-limit are not represented here). In the plot, n, m and fn, respectively, denote the number of cells, the number of mutations and the false negative error rate.

### 4.3 Comparison of PhISCS-BnB against SCITE on simulated data

In a final experiment, we compared PhISCS-BnB against one of the best-known tools for tumor phylogeny reconstruction, SCITE (Jahn et al., 2016), this time with respect to accuracy. As mentioned earlier, SCITE is based on MCMC and as such requires the user to specify the number of iterations, thus indirectly its running time.^7^ As input data, we simulated tumor phylogenies, each with 100 to 300 cells and 100 to 300 mutations, with false negative error rates ranging from 5% to 20%. In each case, 10 distinct trees of tumor evolution were generated, each with 10 subclones. We allowed SCITE to run with 3 restarts, each with a running time (the number of iterations allowed was calculated by dividing this time with the average time per iteration we calculate) 10 times that of PhISCS-BnB on the same input (again, details of the computational platform we used can be found in Supplementary File Section C.1) giving a significant advantage to SCITE over PhISCS-BnB.

For computing the *accuracy* of the inferred tumor phylogenies in comparison to the ground truth, we first used the multi-labeled tree similarity measure (MLTSM) (Karpov et al., 2019) introduced recently. Since MLTSM is a normalized similarity measure, the closer its value to 1.0 implies a higher level of similarity between the inferred tree and the ground truth. The MTLSM between the ground truth trees and the inferred trees is presented in Figure 3. As can be expected, since PhISCS-BnB constructs the optimal tree, the similarity of its output to the ground truth is ∼ 1.0 for all data sets. On the other hand, even though SCITE was given significantly more running time than required by PhISCS-BnB, its output has a relatively low similarity to the ground truth (in the range of [0.2, 0.6]).

**Figure 3:**
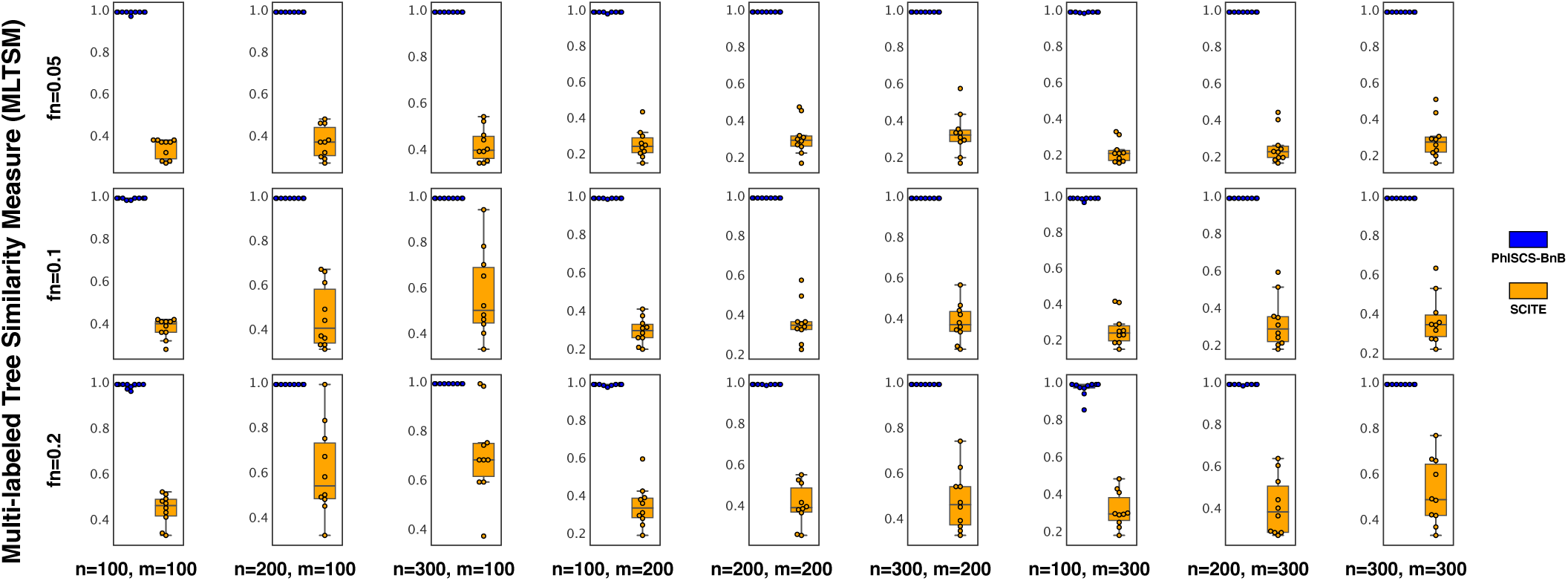
Comparison of PhISCS-BnB with SCITE with respect to the multi-labled tree similarity measure (MLTSM). For each panel, 10 distinct trees of tumor evolution were generated, each with 10 subclones. In the plot, n, m and fn, respectively, denote the number of cells, the number of mutations and the false negative error rate.

We have additionally compared the trees obtained by both SCITE and PhISCS-BnB with respect to other measures as can be seen in Supplementary File Section C.4. With respect to almost every measure, SCITE offers inferior performance in the given time-limit. As a final experiment, we have let SCITE to run on the same datasets for approximately 24 hours. As can be seen in Supplementary File Section C.5, SCITE can then produce trees almost as similar to the ground truth as those obtained by PhISCS-BnB.

## 5 Conclusions

We presented new algorithms and based on them a software package, PhISCS-BnB, to solve the perfect phylogeny problem on noisy single cell (mutation) sequencing data from tumors. On both simulated data and real data from mouse melanoma cell lines, we showed that PhISCS-BnB is one to two orders of magnitude faster than the best available methods, and can solve large instances of practical importance to optimality.

PhISCS-BnB is a branch and bound method, employing a variety of bounding techniques that use either state-of-the-art solvers for classical NP-hard problems such as max-SAT or polynomial time algorithms for 2-SAT and MWM, to prune efficiently and effectively the search tree of solutions. In theoretical computer science, different NP-complete problems are presented as equivalent and reducible to one another (in polynomial time) (Cormen et al., 2009). However, the disproportionate practical importance of a few NP-complete problems, such as max-SAT and the Traveling Salesperson Problem (TSP) (Applegate et al., 2006), has led to high-quality software that can solve instances of these NP-complete problems efficiently and to optimality. Therefore efficient reduction of an NP-complete problem (such as PP) to max-SAT or TSP can leverage existing software to solve the problem much more efficiently. Even if the reduction does not preserve solutions (as per our reductions to 2-SAT or MWM problems), but only gives a bound on solution costs, the reduction can be used within a branch-and-bound framework, as we have done in PhISCS-BnB. This paradigm is widely applicable in bioinformatics in which domain-specific NP-complete problems abound (Gusfield, 2019).

Better understanding of SCS data may lead to better treatments or better strategies for drug development. In current treatment strategies, mutation sequencing data are presented to tumor boards to decide the course of treatment (Mueller et al., 2019). In that clinical context, the rapid availability of phylogenetic trees to identify the tumor subclones could inform treatment. Hence, improving the efficiency of phylogenetic analysis of tumor data, as we have done in PhISCS-BnB, could have a direct impact on clinical treatment decisions.

## Acknowledgements

This work is supported in part by the Intramural Research Program of the National Institutes of Health, National Cancer Institute. This work utilized the computational resources of the NIH HPC Biowulf cluster (http://hpc.nih.gov) and in particular Gurobi (http://www.gurobi.com) to solve some optimization problems. This work was also supported in part by Lilly Endowment, Inc., through its support for the Indiana University Pervasive Technology Institute (Stewart et al., 2017). E.S.A., F.E. and S.C.S. were supported in part by NSF grant AF-1619081 and F.R.M. was supported in part by Indiana U. Grand Challenges Precision Health Initiative.

## Supplemental Material

### A Implementation

We have implemented the algorithm proposed in this paper with pybnb (Hackebeil, 2018) framework. This is a parallel branch-and-bound engine written in Python. It is designed to run on distributed computing architectures, using mpi4py (Dalcin et al., 2011) for fast inter-process communication.

### B Real data analysis

The parental line (P10) and 24 clonal sub-lines (C1-C24) were submitted for exome sequencing to reach 100x coverage. Fastq sequence reads were mapped to the mouse reference genome mm10 with BWA (Li and Durbin, 2009) or Bowtie (Langmead and Salzberg, 2012). Single nucleotide variants (SNV) were identified using samtools mpileup (Li et al., 2009) or GATK HaplotypeCaller (DePristo et al., 2011). Mouse germline single nucleotide polymorphisms (SNPs) were filtered out the Sanger database for variants identified from whole genome sequencing of 36 mouse strains^8^. Variants with a Phred-scaled quality score of <30 were removed. Variants that are present in normal spleen samples (in-house collection) were also removed. Variants were annotated with Annovar (Wang et al., 2010) software to identify non-synonymous mutations.

### C Benchmarking SCITE, PhISCS-I, PhISCS-B and PhISCS-BnB

#### C.1 First platform

Some of the experiments in this work were performed using the *Carbonate*^9^ system, a computer cluster at Indiana University. We used compute nodes from this cluster that are a Lenovo NeXtScale nx360 M5 server equipped with two 12-core Intel Xeon E5-2680 v3 CPUs and four 480 GB solid-state drives. All nodes run Red Hat Enterprise 7.x. We allowed our experiments to use up to 40GB of RAM.

#### C.2 Second platform

Some of the experiments in this work were performed using the *Biowulf* ^10^ system, a computer cluster at National Institutes of Health (NIH). We used compute nodes from this cluster that have Intel E5-2650v2 CPUs.

#### C.3 Running SCITE other options

For SCITE, setting -fd parameter to 0 lead to the segmentation fault. Therefore we set the value of this parameter to 0.0000001. Parameter -e related to the probability of learning noise rates in a given MCMC step was set to 0.2. The full command used to run SCITE is given below:

~~~
scite \
    -i $PATH_TO_INPUT_FILE \
    -names $PATH_TO_GENE_NAME_FILE \
    -n $n \
    -m $m \
    -ad $fn \
    -fd 0.0000001 \
    -e 0.20 \
    -r 3 \
    -l $iterations \
    -o $PATH_TO_OUTPUT > $PATH_TO_OUTPUT.log
~~~

#### C.4 Comparison of PhISCS-BnB against SCITE by other measure

**Figure 1:**
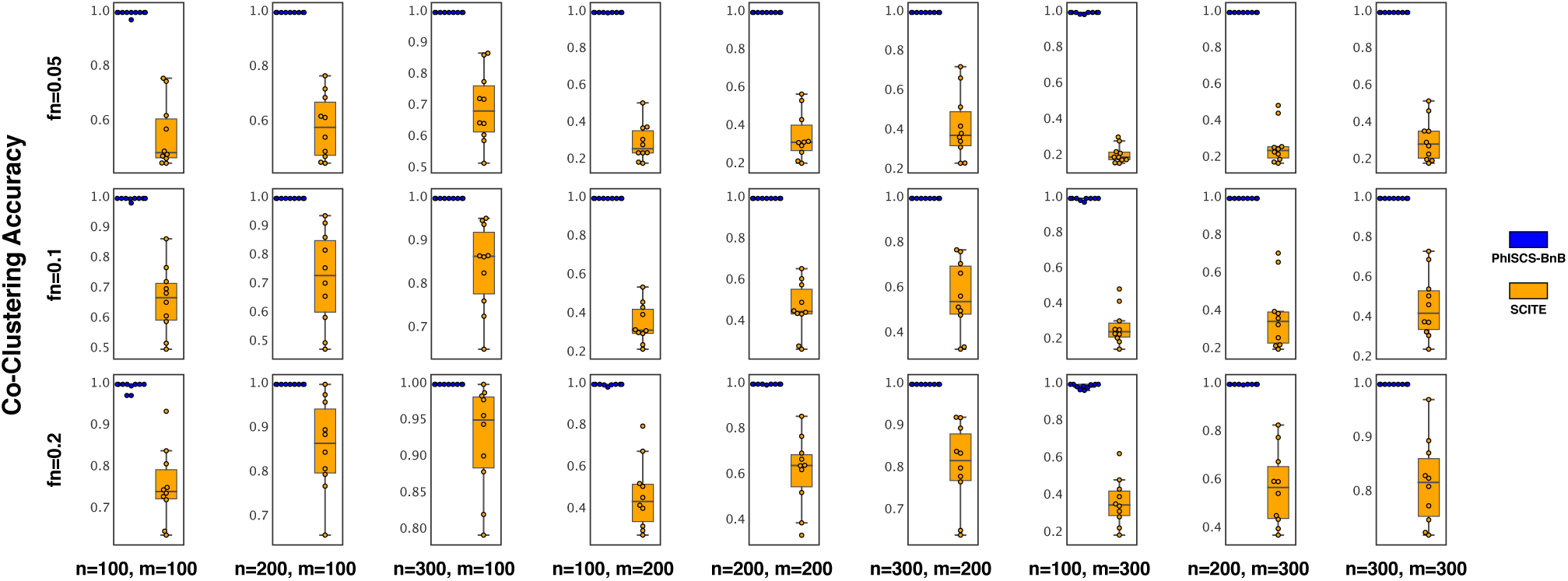
Comparison of PhISCS-BnB with SCITE with respect to the co-clustering accuracy measure. For each panel, 10 distinct trees of tumor evolution were generated, each with 10 subclones. In the plot, n, m and fn, respectively, denote the number of cells, the number of mutations and the false negative error rate. SCITE was allowed to run with 3 restarts, each with a running time, 10 times that of PhISCS-BnB on the same input.

**Figure 2:**
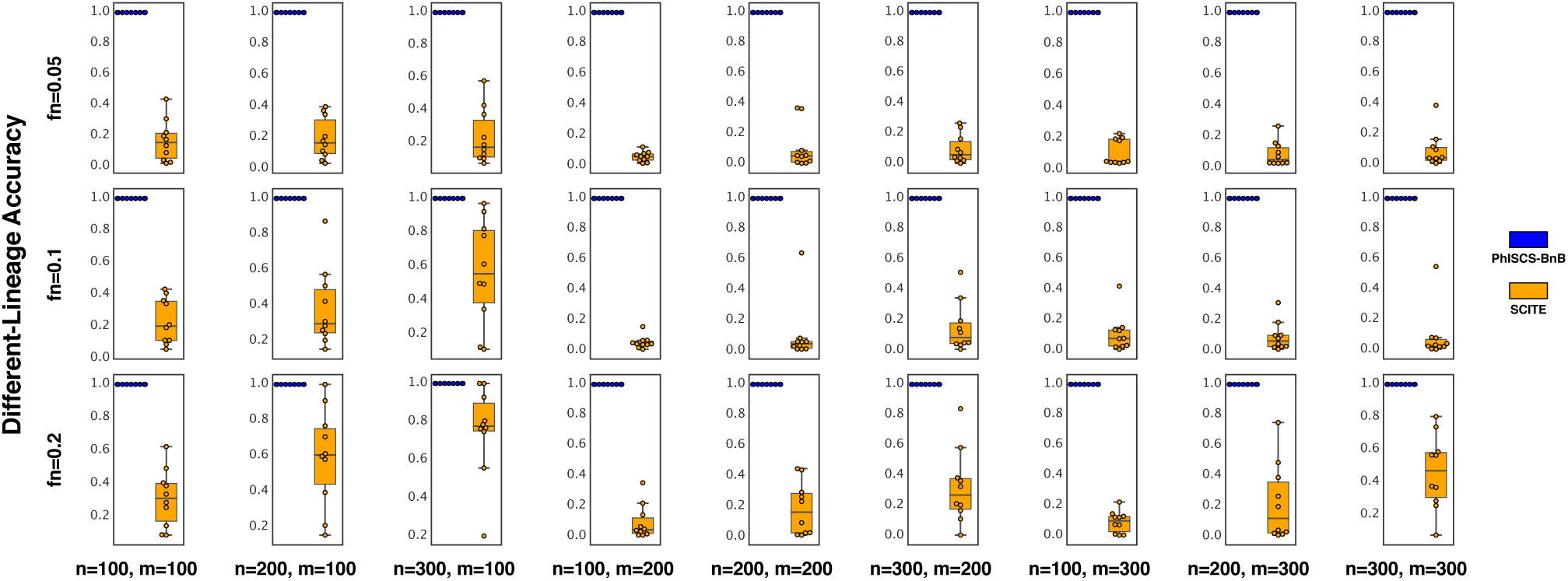
Comparison of PhISCS-BnB with SCITE with respect to the different-lineage accuracy measure. For each panel, 10 distinct trees of tumor evolution were generated, each with 10 subclones. In the plot, n, m and fn, respectively, denote the number of cells, the number of mutations and the false negative error rate. SCITE was allowed to run with 3 restarts, each with a running time, 10 times that of PhISCS-BnB on the same input.

**Figure 3:**
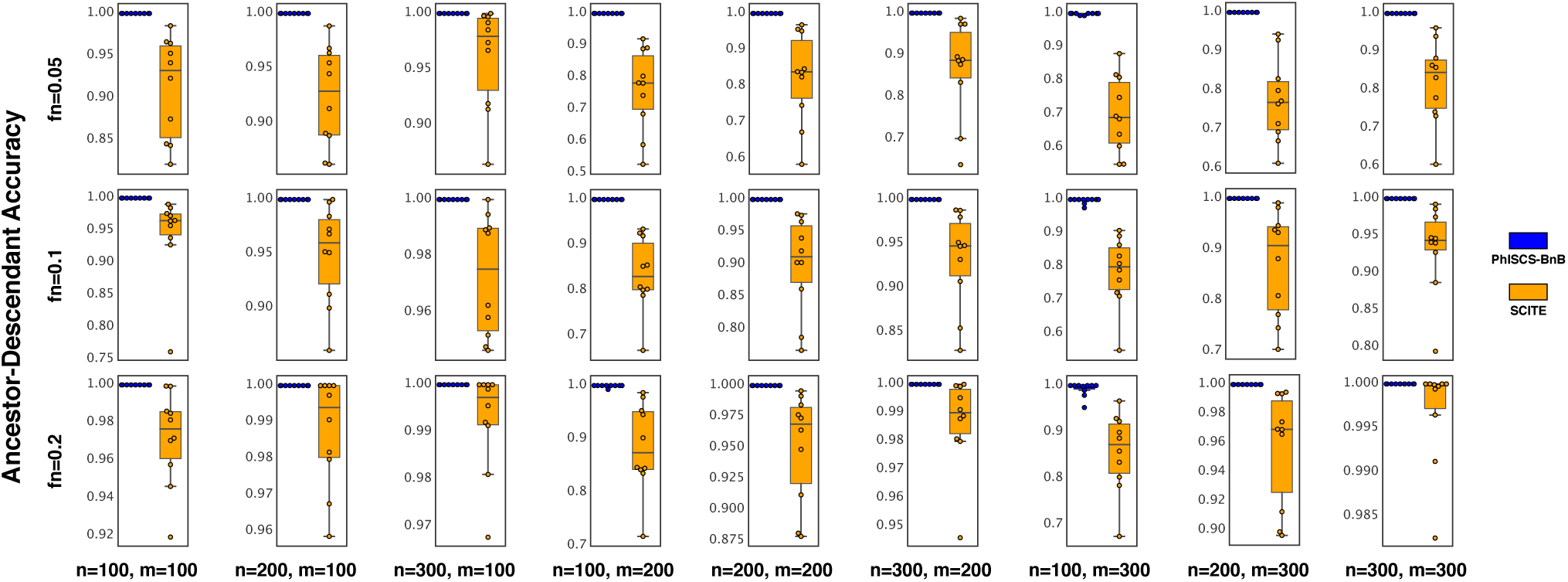
Comparison of PhISCS-BnB with SCITE with respect to the ancestor-descendant accuracy measure. For each panel, 10 distinct trees of tumor evolution were generated, each with 10 subclones. In the plot, n, m and fn, respectively, denote the number of cells, the number of mutations and the false negative error rate. SCITE was allowed to run with 3 restarts, each with a running time, 10 times that of PhISCS-BnB on the same input.

#### C.5 Comparison of PhISCS-BnB against SCITE allowing to run approximately for 24 hours

**Figure 4:**
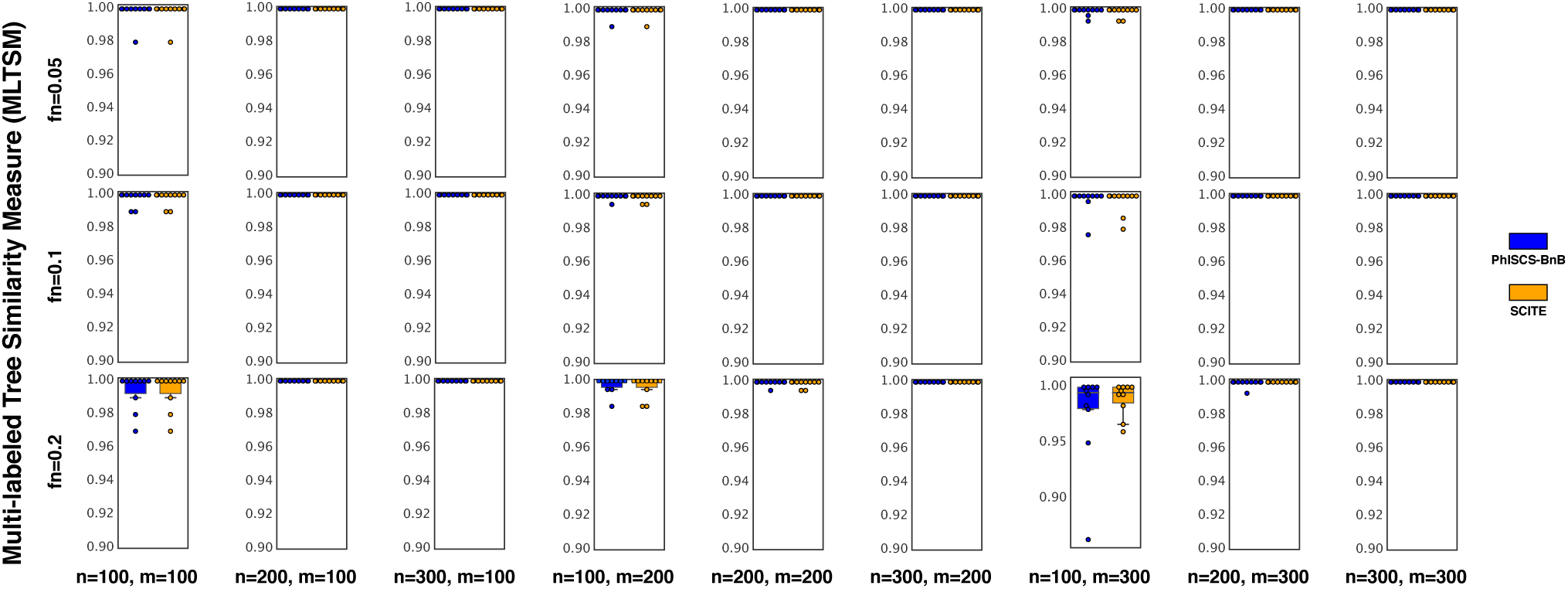
Comparison of PhISCS-BnB with SCITE with respect to the multi-labled tree similarity measure (MLTSM). For each panel, 10 distinct trees of tumor evolution were generated, each with 10 subclones. In the plot, n, m and fn, respectively, denote the number of cells, the number of mutations and the false negative error rate. SCITE was allowed to run approximately for 24 hours.

**Figure 5:**
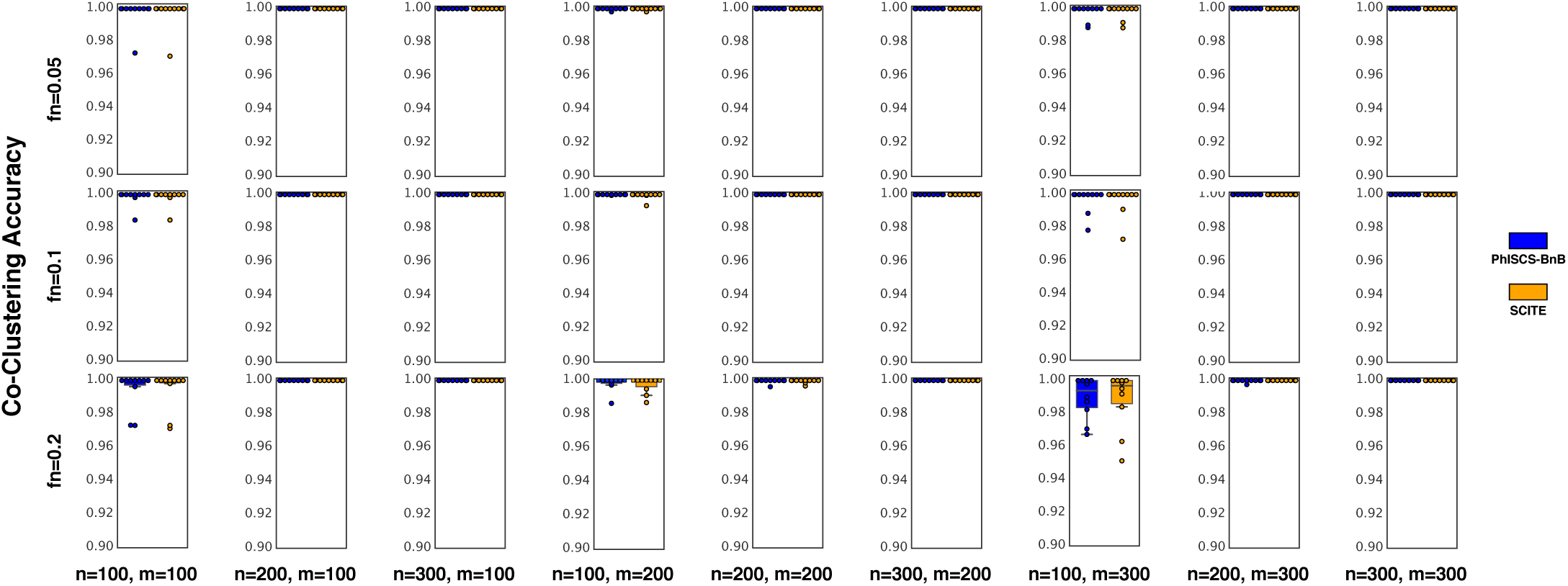
Comparison of PhISCS-BnB with SCITE with respect to the co-clustering accuracy measure. For each panel, 10 distinct trees of tumor evolution were generated, each with 10 subclones. In the plot, n, m and fn, respectively, denote the number of cells, the number of mutations and the false negative error rate. SCITE was allowed to run approximately for 24 hours.

**Figure 6:**
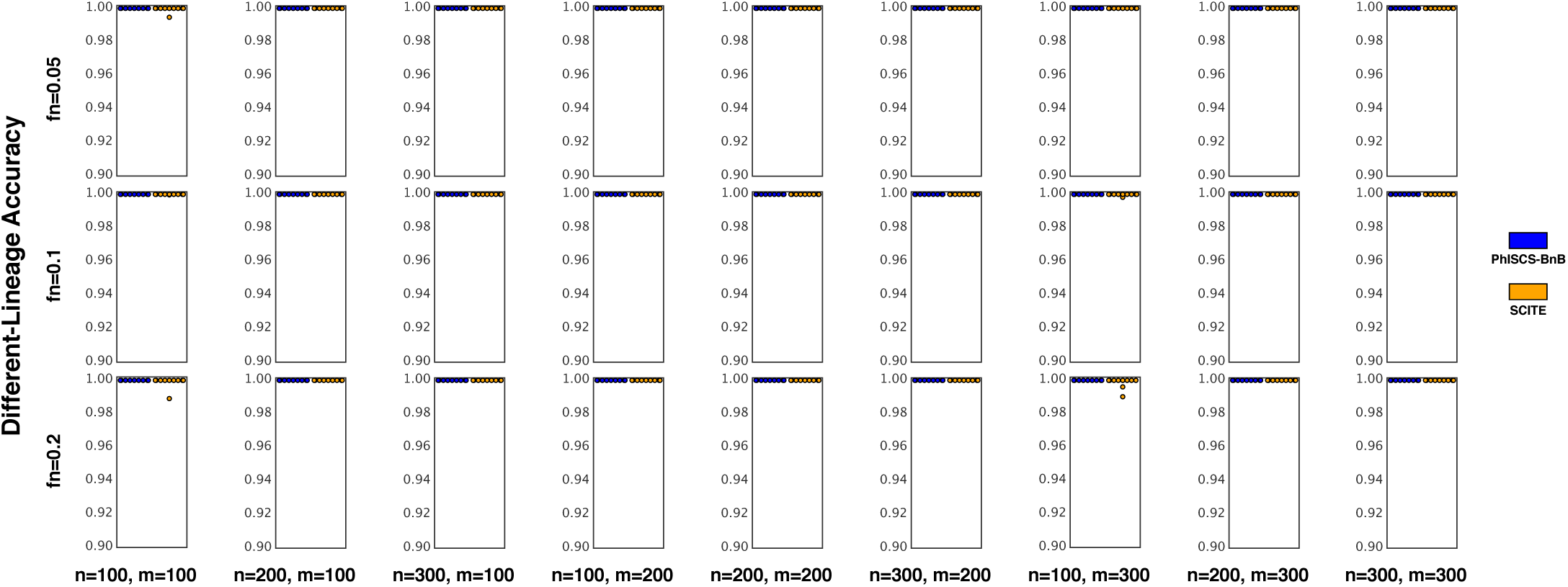
Comparison of PhISCS-BnB with SCITE with respect to the different-lineage accuracy measure. For each panel, 10 distinct trees of tumor evolution were generated, each with 10 subclones. In the plot, n, m and fn, respectively, denote the number of cells, the number of mutations and the false negative error rate. SCITE was allowed to run approximately for 24 hours.

**Figure 7:**
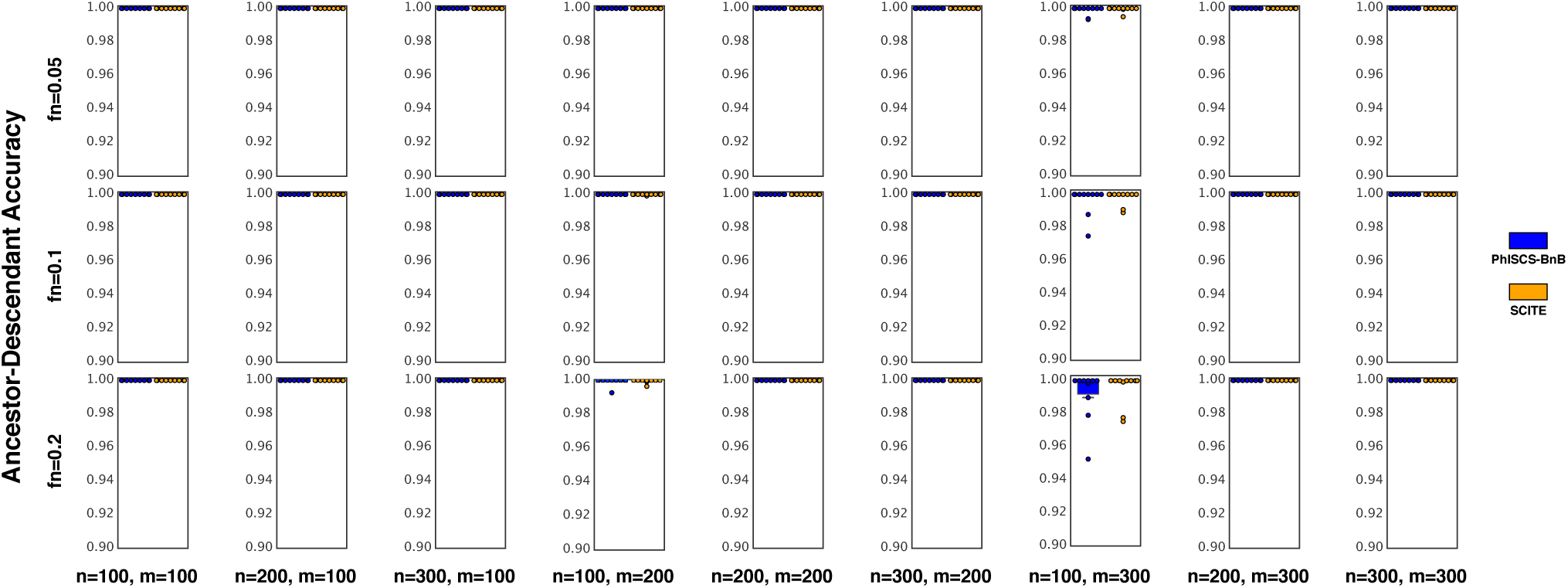
Comparison of PhISCS-BnB with SCITE with respect to the ancestor-descendant accuracy measure. For each panel, 10 distinct trees of tumor evolution were generated, each with 10 subclones. In the plot, n, m and fn, respectively, denote the number of cells, the number of mutations and the false negative error rate. SCITE was allowed to run approximately for 24 hours.

### D Packages used in this work

- SciPy (Virtanen et al., 2019)
- NumPy (Oliphant, 2006)
- pandas (McKinney, 2010)
- Matplotlib (Hunter, 2007)
- OR-Tools (Perron and Furnon, 2019)
- tqdm (da Costa-Luis, 2019)
- NetworkX (Hagberg et al., 2008)

## Footnotes

1 A final noise source is the *doublets*, the technical artifacts of two (or rarely more) cells with heterogeneous mutation profiles treated and sequenced as a single cell. Since there are a number of preprocessing techniques such as (Roth et al., 2016) to detect and eliminate doublets fairly well we will not focus on doublets as a source of noise in this paper.

2 SiFit also allows for deletion events and loss of heterozygosity.

3 One exception is ScisTree (Wu, 2019) which is reported to be faster but is a heuristic approach with no optimality guarantees.

4 In general both false negative and false positives (respectively, 1 read as 0 and 0 read as 1) happen with distinct probabilities. The qualitative difference in these probabilities is due to the sequencing technology in use and thresholding rules employed in establishing *I*. As is well known, the false positive rate is typically much lower than the false negative rate. In fact, in emerging data, e.g. from SClS experiments, the false positive rate approaches zero and thus can be ignored. As a result we focus only on false negatives and our proposed algorithm and its sub-routines make use of this assumption.

5 If there are multiple pairs of columns involved in violations we impose an ordering on them and pick a pair according to this order; thus, we are always considering a single column pair.

6 Note that we used the compact formulation that is mentioned in Section 3.2.2 to run PhISCS-BnB on the simulated data but not on real data.

7 The approximation of the time that SCITE takes per iteration for a given input matrix was calculated by running it 10 times, each with 20000 iterations with 1 restart and then taking the average running time per iteration in this set of runs.

8 ftp://ftp-mouse.sanger.ac.uk/current_snps/mgp.v5.merged.snps_all.dbSNP142.vcf.gz

9 https://kb.iu.edu/d/aolp#overview

10 https://hpc.nih.gov/systems/

